# Population dynamics and ranging behaviours of provisioned silvered langur (*Trachypithecus cristatus*) in Peninsular Malaysia

**DOI:** 10.1101/2020.12.16.423156

**Authors:** Norlinda Mohd-Daut, Ikki Matsuda, Kamaruddin Zainul Abidin, Badrul Munir Md-Zain

## Abstract

Tourists are attracted to the Bukit Melawati Kuala Selangor (BMKS) of Peninsular Malaysia, a small hill park, both for its status as a historical site and the free-ranging silvered langurs (*Trachypithecus cristatus*) that come for provisioning. We assessed the population trends and group sizes of *T. cristatus* over 10 years in the BMKS and examined their ranging patterns. Comparisons of observed populations between 2005 (190 individuals) and 2017 (193 individuals) revealed the stable demography and group sizes of the six *T. cristatus* groups in the BMKS. Based on a total of 185 location points of the six groups in 2017, their mean ranging area was 3.6 ha with a range of 0.86 to 6.93 ha with extensive spatial overlap. We also found a significant positive relationship between the six groups’ ranges and group sizes in 2017. Additionally, qualitative ecological comparisons with a previous study on *T. cristatus* in 1965 (before provisioning) suggest that the artificial food supply in the study area could modify the population dynamics and socioecology of *T. cristatus*. The modifications might alter their range size and territoriality in the BMKS. Overall, we found that provisioning had negative effects on the ecology of *T. cristatus* in the BMKS. Therefore, modifying management policies, such as banning feeding and implementing educational programs, may contribute to their proper conservation.

## Introduction

The monitoring of animal populations is essential for developing effective management plans for long-term conservation (Wich and Marshall 2016). Primates are long-lived mammals with slow life cycles (Charnov and Berrigan 2005; Ross 1998), so they generally respond slowly to dramatic environmental changes and human interference (Chapman et al. 2006). Therefore, long-term population monitoring data is indispensable for understanding their population dynamics (e.g. Chapman et al. 2007; Matsuda et al. 2020). Knowledge of a species’ ranging behaviour could also contribute to a better understanding of its behavioural ecology and habitat requirements, leading to effective conservation strategy designs (Nathan et al. 2008). Among various environmental factors, food distribution and abundance are significant influences on population characteristics, such as demography and density (e.g. Bernard et al. 2019; Chapman et al. 2017; Hanya et al. 2004; Marshall 2010), and ranging in many primate species (e.g. Ampeng and Md-Zain 2012; Clutton-Brock 1975; Green et al. 2020; Hanya et al. 2020; Matsuda et al. 2009; Olupot et al. 1994; Raemaekers 1980; Zhou et al. 2007). Access to more energy-rich foods generally leads to higher birth rates or shorter interbirth intervals (Ross 1998), potentially inducing larger group sizes and reducing the per capita foraging success via more scramble competition for food (Chapman and Chapman 2000; Janson 1988). In theory, this should be associated with larger groups of primates traveling greater distances and having larger home ranges to meet their nutritional requirements (Makwana 1978; Takasaki 1981).

As the availability and quality of resources are closely related to demography and ranging patterns of primates, both direct (e.g. being fed by tourists) and indirect (e.g. garbage eating) provisioning have been reported to influence the demography (Altmann and Alberts 2003; Borries et al. 2001; Mori 1979; Sugiyama and Ohsawa 1982; Takahata 1980) and ranging patterns of primates (Altmann and Muruthi 1988; Asquith 1989; Kurita et al. 2008; Md-Zain et al. 2010; Ruslin et al. 2019; Sha and Hanya 2013). When primates are provisioned, their daily travel distance and total ranging area generally decreases (Altmann and Muruthi 1988; José-Domínguez et al. 2015; Sengupta et al. 2015) since food is provided by humans to individuals or groups in specific areas. However, it should be noted that provisioning nutrient-rich foods generally lead to higher birth rates or shorter interbirth interval (Rothman and Bryer 2019); subsequently, group sizes will increase with increasing daily travel distances, as this is a general effect regardless of provisioning.

As a result of provisioning, animals are often over-habituated (Russon and Wallis 2014b). Therefore, provisioning can trigger human-animal conflict, which is of great concern to conservationists and wildlife managers. For example, several reported conflicts have involved situations such as monkeys attacking and injuring tourists, possibly infecting tourists and vice versa, and damage to the natural habitat (e.g. Brennan et al. 1985; Fuentes et al. 2007; Kurita 2014; Md-Zain et al. 2011; Palacios et al. 2011; Sha et al. 2009). Consequently, ecotourism that includes free-ranging primates, especially concerning provisioning, should be implemented with caution.

Silvered langurs (*Trachypithecus cristatus*), classed as vulnerable by the IUCN and widely distributed in Southeast Asia (Meijaard and Nijman 2015), are foregut-fermenting colobines (Chivers 1994; Matsuda et al. 2019). The ecological knowledge of *T. cristatus* is limited; however, according to a preliminary study (Hock and Sasekumar 1979), they are predominantly folivorous, with leaves comprising over 90% of their diet.

Tourists are attracted to the Bukit Melawati Kuala Selangor (BMKS), a small hill park in Peninsular Malaysia, due to its status as a historical site, its beautiful offshore scenery and the presence of free-ranging habituated *T. cristatus* as a result of tourist provisioning (e.g. feeding the animals bread or cookies) (Mohd-Daut 2020). Before provisioning, the *T. cristatus* ranging area was strictly territorial, and the ranges of adjacent groups overlapped by only a few trees in the BMKS (Bernstein 1968). Their home range has been estimated to be 20 ha in size, with 350 m as the mean daily path length (Yeager and Kool 2000). Although it is unclear when tourist provisioning started, the *T. cristatus* in the site appear to be over-habituated as early as the 2000s. They have been observed approaching visitors to beg for food, staring at and touching tourists and stealing food from tourists (Mohd-Daut and Md-Zain 2021, and see Fig. 1); however, the langurs do not initiate aggressive contact. Only one study (Hambali et al. 2016) reported the non-territorial ranging behaviour of one *T. cristatus* group in the site (home range: 1.0 ha; daily path length: 10–200 m), though the information on the ranging patterns in other adjacent groups was not available. Additionally, we have almost no empirical information about their ecology and behaviours in relation to the effects of provisioning in the BMKS based on long-term monitoring.

**Figure 1.** Over-habituated *T. cristatus* in BMKS. They have been approaching tourists to beg for food, to stare at and touch them and to steal their food (photos were taken by NMD and BMM). * This figure including some photographs of the study area cannot be uploaded to bioRxiv due to their policy. Noted that the figure is available from the corresponding authors by reasonable request.

In the present study, to gain a complete picture of the demography and ranging patterns of the provisioned groups of *T. cristatus* in the BMKS, which possibly contribute to the development of their effective management plans, we sought to (1) evaluate population changes between 2005 and 2017 and (2) estimate the range size of all groups at the site. The effects of provisioning on their demography and ranging behaviour are discussed by qualitatively comparing records from a previous study, conducted before tourist provisioning (Bernstein 1968).

## Materials and Methods

We observed six groups of silvered langurs *T. cristatus* from March to April 2005 and from October to December 2017 in a part of the BMKS (3°20′ 26.36 N, 101°14′42.46 E), located approximately 62 km from Kuala Lumpur, Malaysia (Hambali et al. 2016). The study area covers ca. 22.5 ha, surrounded by the Kuala Selangor Nature Park, a mangrove forest along the coastline that includes a residential area. Historical structures and monkey habitats in this open area attract visitors and make the BMKS a renowned tourism area in Selangor. Although fragmented, the study area still harbours plants that are important resources in the *T. cristatus* diet, including both natural and ornamental plants (Mohd-Daut and Md-Zain 2021). The mean minimum and maximum temperatures were approximately 26.5°C and 29.8°C, respectively, and total precipitation at the site was 1,505.5 mm (Metereology Department of Malaysia).

We observed the groups with one adult male, multiple adult females and immatures, which were well-habituated to humans. We estimated each individual’s approximate age using age/sex categories based on the criteria of Aggimarangsee (2002), Bernstein (1968) and Harding (2010), with some modifications. In 2005, we identified all *T. cristatus* adult, subadult and juvenile individuals in the study groups based on 1) the shape of the face, especially that distinguished by the curve of the cheek fur and if their faces are □tapered□ (i.e. wider on the angle of the head and smaller on the chin) or □dented□ (i.e. the middle nasal meatus is much lower than the tip of the nose); 2) the shape of the nose and eyes, which usually had characteristics resembling those of the original parent in the group and 3) any visible evidence of defects or injuries, such as scars or limb defects. Although all individuals identified in 2005 were not recognised in 2017, some females in all groups (mean = 2.7 ± 2.7 individuals per group; range = 1–8 individuals) with distinctive features, such as scars and limb defects, were recognisable even in 2017. Notably, all adult males were replaced from 2005 to 2017. Thus, in 2017, the recognisable individuals were adult females, and the groups including such females were found in similar areas to those used by the groups they were member of in 2005. In addition to the fact that *T. cristatus* appears to be a species characterised by female philopatry and male-biased dispersal (Wolf and Fleagle 1977), we considered this as evidence that these were the same groups. It is unlikely, but even if the groups identified in 2005 and 2017 were different, this would not affect the overall results of this study, especially in the comparison of population size and mean group size.

At 0800 h each day, we arrived at the study site and followed the groups until they selected evening sleeping trees and started sleeping, usually at 1900 h. We completed 520 and 1,023 hr observations in 2005 and 2017, respectively. Once-daily, we checked the numbers, age class, and sex of group members. On the first day of each observation week, we attempted to record the location points for all study groups at the site using Global Positioning Systems (GPS). The location data was typically recorded once a week. Of the six groups, five were constantly observed in the study area, although one group (group F) was infrequently found. We recorded 185 location points for the six *T. cristatus* groups, with 14 to 34 points per group. We transferred GPS coordinates in decimal degrees and projected them into meters (WGS 1984 UTM zone 48N) using the projection tool in ArcMap version 10.5. Biotas 2.0 Alpha was used to calculate and investigate the ranging area. The most common method, namely Minimum Convex Polygon (MCP) 100% (Boitani and Fuller 2000), was used to analyse each group’s home range. Based on the home range areas for each group (calculated by MCP), we also calculated the amount of overlap between the group areas. Google Earth was used to map and estimate the size of the land area used in the study. Areas used by the study groups were divided into three categories: (i) vegetated areas, including all areas covered by trees; (ii) grassland and open areas; and (iii) building/construction areas.

We compared the group size of six one-male-multi-female groups between 2005 and 2017, using a Wilcoxon rank-sum test (Wilcoxon 1945). We examined the relationship between home range size and the six groups’ sizes in 2017, using linear regression analysis. The relationship between the total area size of vegetated patches within each group’s ranging areas and their group size was also tested using linear regression analysis. We used the Bonferroni correction to correct for multiple comparisons of those linear regression analyses (P = 0.05/N: N is the number of comparisons made, i.e. 0.025). The significance level was set to α = 0.05. We report means with standard deviation for all results. All statistical analyses were performed in the R statistical computing environment version 4.02 (R-Core-Development-Team 2019).

## Results

### Population comparison between 2005 and 2017

In 2005, we counted 190 *T. cristatus* in six groups with one male and multiple females at the study site. The population size was almost the same in 2017 (193), though we found an additional all-male group (Table 1). The mean size of groups with one male and multiple females was 31.7 ± 16.5 and 31.5 ± 12.4 individuals in 2005 and 2017, respectively, and the size was not significantly different between the years (W = 17; *p*-value: 0.916). We found that, out of the six one-male-multi-female groups, two groups had more members (i.e. A and F), and two groups maintained their group size (i.e. B and E). In contrast, the other two groups had fewer members (i.e. C and D). Groups A and F experienced an increase of 10 and 2 members, respectively. On the other hand, Group C experienced a decrease of 3 members in the total group size of 24 individuals. Meanwhile, Group D, which consisted of 46 individuals, decreased by 10 members.

**Table 1.** Composition of groups of *Trachypithecus cristatus*, with several one-male-multifemale groups (OMG) and an all-male group (AMG) in 2005 and 2017.

| Group ID | Group type | Year | Age and sex categories |  |  |  |  |  | Total |
| --- | --- | --- | --- | --- | --- | --- | --- | --- | --- |
|  |  |  | Adult male | Adult female | Subadult male | Subadult female | Juvenile | Infant |  |
| A | OMG | 2005 | 1 | 12 | 0 | 1 | 3 | 2 | 19 |
|  |  | 2017 | 1 | 14 | 1 | 4 | 7 | 2 | 29 |
| B | OMG | 2005 | 1 | 22 | 3 | 2 | 2 | 3 | 33 |
|  |  | 2017 | 1 | 21 | 2 | 2 | 4 | 3 | 33 |
| C | OMG | 2005 | 1 | 13 | 4 | 3 | 3 | 3 | 27 |
|  |  | 2017 | 1 | 18 | 1 | 1 | 1 | 2 | 24 |
| D | OMG | 2005 | 1 | 8 | 7 | 11 | 24 | 5 | 56 |
|  |  | 2017 | 1 | 32 | 0 | 7 | 2 | 4 | 46 |
| E | OMG | 2005 | 1 | 28 | 1 | 1 | 7 | 6 | 44 |
|  |  | 2017 | 1 | 33 | 2 | 4 | 1 | 3 | 44 |
| F | OMG | 2005 | 1 | 7 | 0 | 3 | 0 | 0 | 11 |
|  |  | 2017 | 1 | 5 | 1 | 2 | 2 | 2 | 13 |
| G | AMG | 2005 | - | - | - | - | - | - | - |
|  |  | 2017 | 4 | 0 | 0 | 0 | 0 | 0 | 4 |

### Ranging area in 2017

Based on 185 location points of six one-male-multi-female groups, their mean ranging size was 3.6 ± 2.08 ha with a range of 0.86 to 6.93 ha (Table 2 and Fig. 2). The size of the area calculated by MCP using the cumulative location points in each group suggests that we have not fully recorded the total area occupied by the study groups as the area size did not fully reach a plateau for each group (Fig. 3). Nonetheless, within their ranging areas, the most dominant habitat type was the vegetated area (54.8 ± 7.34 ha), followed by the grass/open area (32.7 ± 5.35) and the building/construction area (12.6 ± 3.99), though the percentage of each habitat type was different across the six groups (Table 3).

**Table 2.**
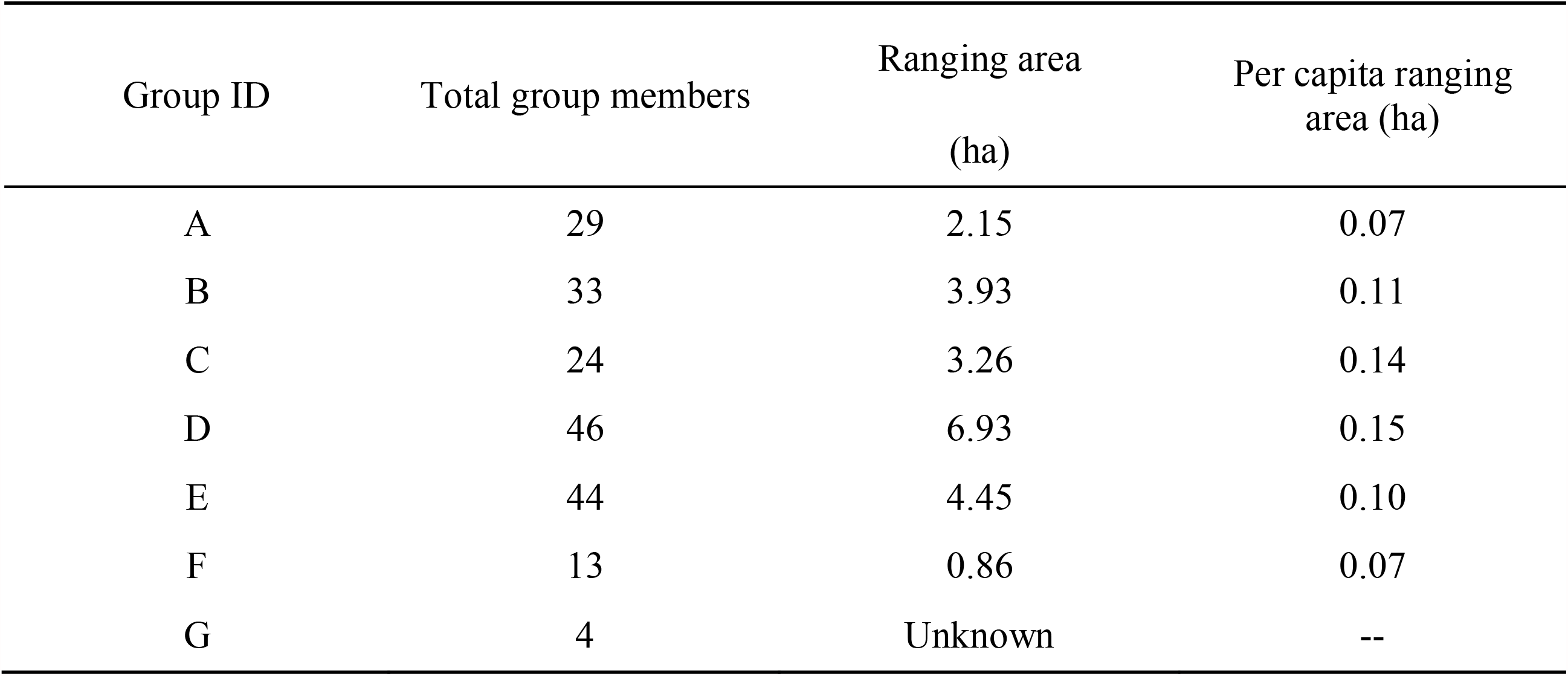
Range area and per capita ranging area of the study *T. cristatus* groups

**Table 3.** Percentage of the land used for each *T. cristatus* group.

| Group | A | B | C | D | E | F |
| --- | --- | --- | --- | --- | --- | --- |
| Vegetated area | 47.04 | 61.10 | 47.42 | 61.54 | 61.55 | 49.87 |
| Grass and open area | 37.72 | 29.10 | 41.01 | 28.95 | 28.51 | 30.64 |
| Building/construction area | 15.24 | 9.89 | 11.56 | 9.51 | 9.94 | 19.49 |

**Figure 2.**
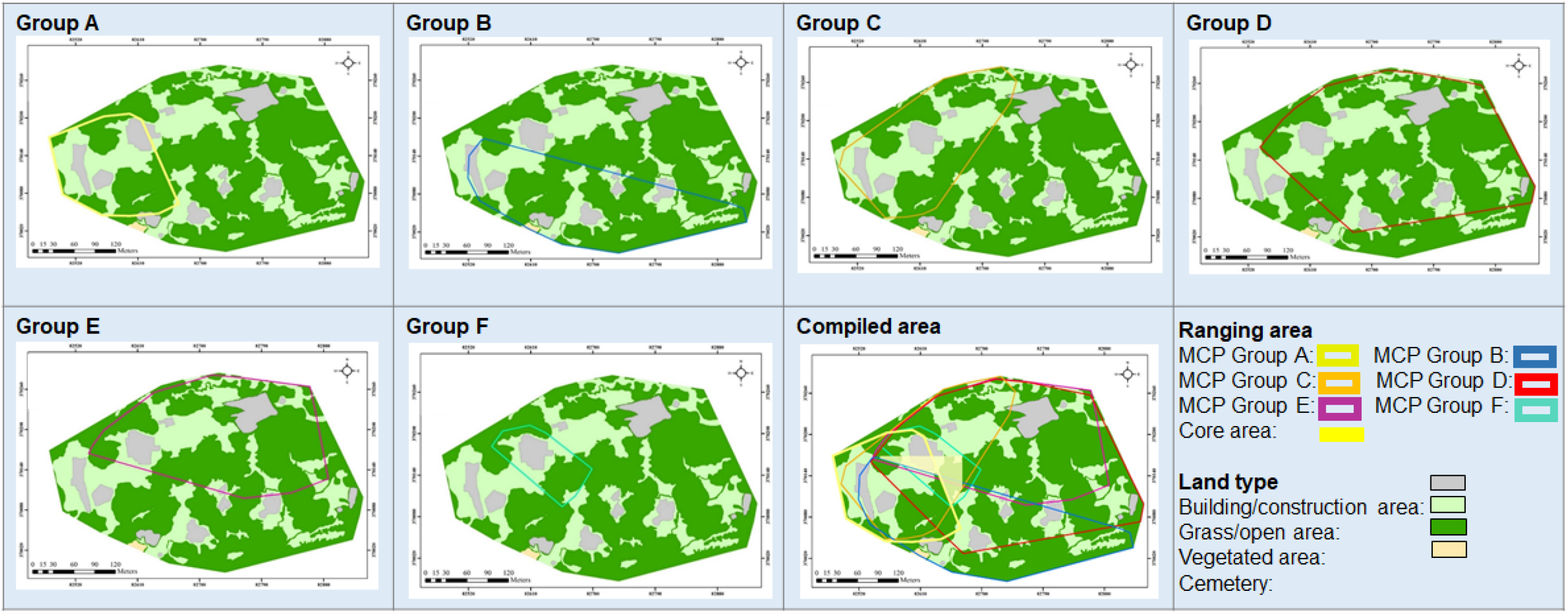
Ranging area of each *T. cristatus* group (Group A–E) with compiled ranging areas, i.e. the core area used by all the six groups in 2017.

**Figure 3.**
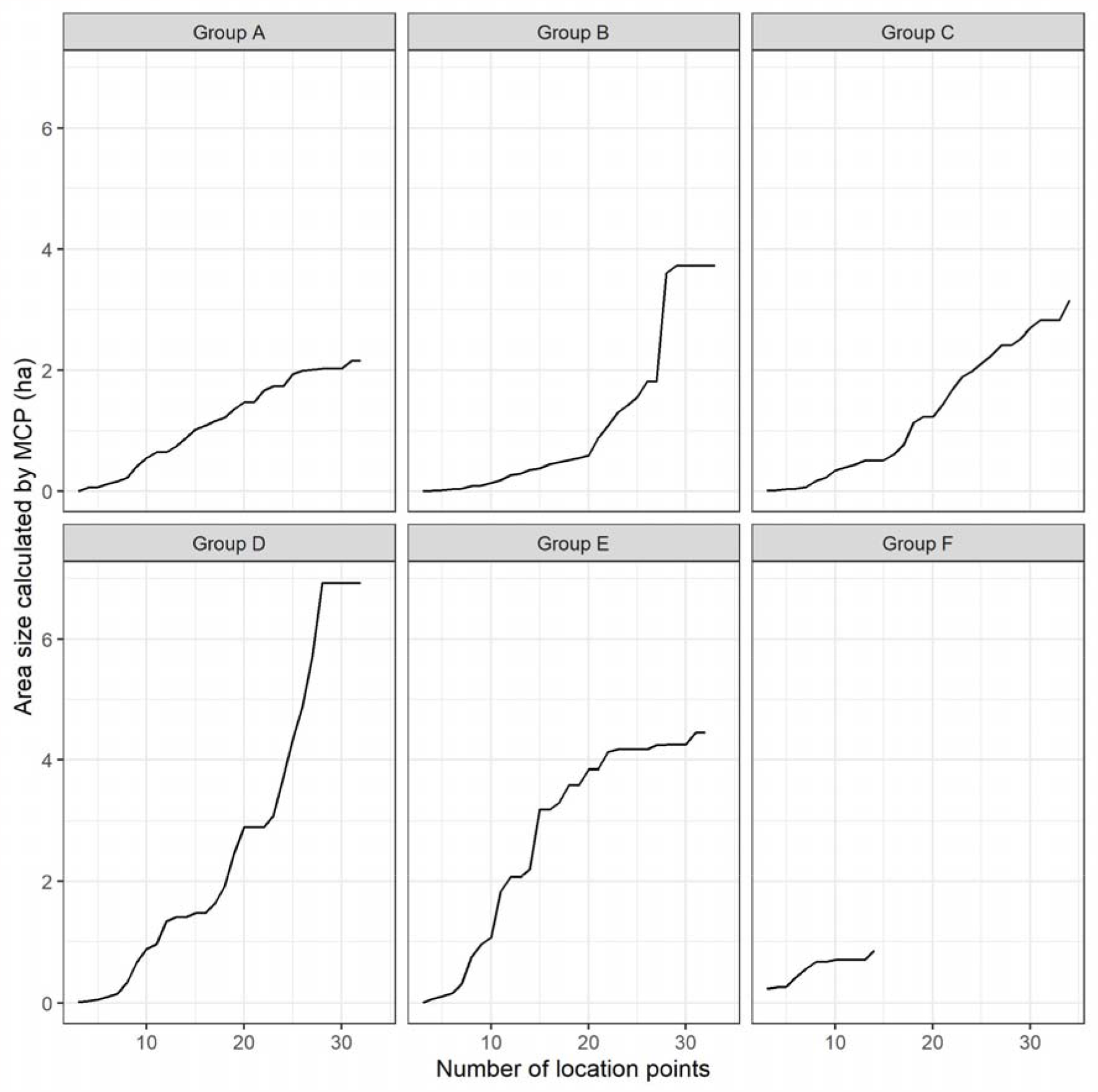
Area size calculated by MCP based on the cumulative location points in each study group.

There was range area overlap among the six one-male-multi-female groups. On average, Group A shared 44.7% ± 26.5% (range: 19.0%–80.2%), Group B shared 22.0% ± 21.7% (3.0%–48.2%), Group C shared 47.5% ± 20.4% (24.7%–71.2%), Group D shared 34.1% ± 21.4% (12.4%–63.3%), Group E shared 39.5% ± 42.5% (2.7%–98.6%) and Group F shared 74.6% ± 34.2% (24.7%–99.6%) of their home range with five other groups. In summary, the mean overlapping area of the six groups based on all dyadic combinations was 41.8% ± 28.9% (1.28% ± 1.08 ha), with a range of 3.0%–99.6% (0.12–4.39 ha) (Table 4). The core area, which was used by all six groups, was 0.33 ha (Fig. 2).

**Table 4.** Asymmetrical matrix of the spatial overlap in percentage data point between the six one-male multifemale groups monitored in the study area. Horizontal rows correspond to the percentage of overlapping between two different groups.

| Group ID | A | B | C | D | E | F |
| --- | --- | --- | --- | --- | --- | --- |
| A |  | 61.1 | 80.2 | 43.4 | 19.0 | 19.8 |
| B | 33.4 |  | 31.5 | 48.2 | 3.0 | 5.4 |
| C | 52.9 | 38.0 |  | 71.2 | 56.1 | 24.7 |
| D | 13.5 | 27.4 | 33.5 |  | 63.3 | 12.4 |
| E | 9.2 | 2.7 | 41.1 | 98.6 |  | 15.6 |
| F | 49.5 | 24.7 | 93.5 | 99.6 | 80.6 |  |

The six groups’ per capita ranging size differed across the groups with a range of 0.07 to 0.15 ha (Table 2). There was a significant positive relationship between their ranging area and group sizes (Fig. 4; *R*^*2*^: 0.81; corrected *p*-value: 0.030). There was also a significant positive relationship between the vegetated area within each group’s range and group sizes (Fig. 4; *R*^*2*^: 0.82; corrected *p*-value: 0.025).

**Figure 4.**
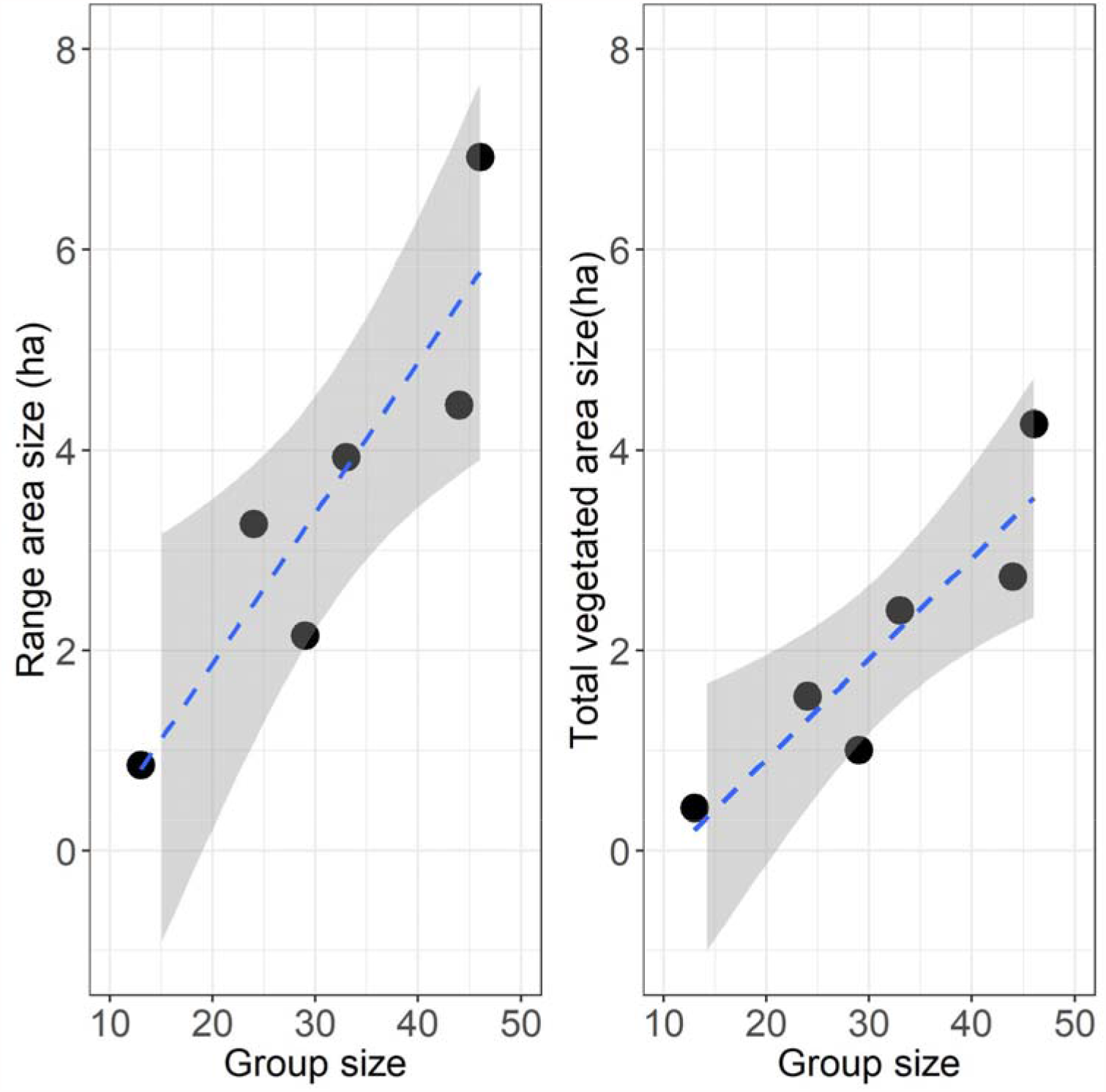
Relationships between the ranging area in each group (left) and the total vegetated area within each group’s ranging area with group size (the number of group members in each group). The blue dotted lines and shaded areas represent the linear regression lines for the observed samples and their 95% confidence interval ranges, respectively.

## Discussion

The population size of *T. cristatus* in the BMKS study area was almost the same in 2005 and 2017 (190 and 193 individuals, respectively), despite the addition of an all-male group, indicating that their numbers have remained stable over the decade. Group sizes in 2017 (27.6) were smaller than in 2005 (31.7), but it was not a significant difference. Although provisioning impacts primate demography in general (Rothman and Bryer 2019), we could not detect significant changes in population or group sizes in the study site, indicating that the population is at equilibrium, i.e. births and immigrations are balanced out by deaths or dispersals. In other words, this result suggests that a constant supply of high nutrient artificial foods provided by tourists in the 2000s (Md-Zain et al. 2009) may have had a minimal effect on the demography of the *T. cristatus*. However, it should be noted that the current information is too limited to comprehensively evaluate the provisioning effects, and a further study focusing on the nutritional intake (in terms of calories, protein, and fiber) of *T. cristatus* in the BMKS is necessary. On the other hand, the density of *T. cristatus* using the area in 2017 was unusually higher (ca. 858/km^2^) than those of *Trachypithecus* spp. in other study sites (Newton and Dunbar 1994): mean value = 101/km^2^; range = 2.1–345/km^2^. As the density of primates is generally influenced by provisioning and/or habitat quality (e.g. Bernard et al. 2019; Chapman et al. 2017; Hanya et al. 2004; Marshall 2010), the fact that the density at our study site is extremely high does imply that provisioning previously results in a high birth rate in our study area. However, as the population is no longer increasing over the decade, mortality (or dispersal from the area) keeps the population size constant.

Although the demography was stable even under the provisioning condition over the decade, it is reasonable to expect the *T. cristatus* population size to increase due to the effects of artificial foods provided by tourists in the 2000s but not in the 1960s in the BMKS (Bernstein 1968). In 1965, the six *T. cristatus* groups, including at least 160 individuals, were reported (Bernstein 1968). Based on these figures, their population size would have increased by ca. 30 individuals (19%) from the 1960s to the 2000s, suggesting that there has been an impact of provisioning on the population size of *T. cristatus*. However, since we only conducted the population survey in a part of the BMKS that tourists can access, additional groups might inhabit the forested area. A population survey of *T. cristatus* in the entire BMKS area would allow us to better conclude how tourist provisioning has influenced the demography over the last five decades.

The artificial food supply in the study area may modify the socioecology of *T. cristatus*, altering their range size and territoriality in the BMKS. The study group ranging areas in 2017 (mean of the six groups = 3.6 ha) were much smaller than the 1965 ranging areas (20 ha: Bernstein 1968; Yeager and Kool 2000), although we cannot deny the possibility that this effect may have been an artefact of the small sample sizes, in terms of the number of location data points for each study group. Nonetheless, it should be noted that MCP, used in this study to calculate the home range sizes, is a more suitable measure of home range area, especially for a data set with fewer location points (e.g. Grueter et al. 2009; Robbins 2003). Furthermore, as the study groups sometimes moved out of the area and briefly inhabited the forested area (NMD personal observations), the real sizes of the study group home ranges might be underestimated. However, we observed that the *T. cristatus* groups mostly spent their time in the BMKS study area and even slept there at night, which would minimize errors in estimating their home range values. It may not be necessary for *T. cristatus* to forage larger areas to satisfy their nutrient requirements as they could easily obtain energy-rich artificial foods in the study area. Such behavioural changes reportedly led to decreases in ranging areas or daily path lengths in other provisioned primate species, e.g. pig-tailed macaques (José-Domínguez et al. 2015), rhesus macaque (Sengupta et al. 2015), and baboons (Altmann and Muruthi 1988). Thus, because no artificial or feeding maintenance occurred in 1965 (Bernstein 1968), the observed change of the *T. cristatus* range sizes in our study area may be influenced by recent tourist provisioning.

Additionally, the six *T. cristatus* group ranging areas were more concentrated to our study area where human food was accessible and abundant in 2017 than those in 1965 (Bernstein 1968). Thus, the population density in 2017 would be more biased to such a specific tourist area of the BMKS. This bias may explain why the six *T. cristatus* groups in 1965 ranged larger areas and were more territorial, with less spatial overlap, among the groups (Bernstein 1968) compared to those in 2017. In the 1960s, the population might have been near or above carrying capacity, and groups would be territorial at high densities in the BMKS. However, when people started provisioning with high nutrient foods, the population might have dipped below carrying capacity, and the groups would be non-territorial. This explanation also fits the hypothesized theoretical relationship among home range overlap, individual density, and carrying capacity, according to Yeager and Kool’s (2000) description of primate territorial behaviours.

Although the ranging behaviours of the study groups would be influenced by the provisioning (i.e. decreased ranging area), there was a significant positive relationship between ranging areas and group sizes of the six *T. cristatus* groups. Theoretically, increases in group size should reduce per capita foraging success (Chapman and Chapman 2000; Janson 1988), i.e. larger groups generally travel greater distances with a larger home range area (e.g. Makwana 1978; Takasaki 1981). We also found a significant positive relationship between vegetated areas within the ranging area of each group and group size, suggesting that provisioned foods may not be sufficient to meet their nutrient requirements. Thus, the larger *T. cristatus* groups would still need to forage natural foods within the larger vegetated areas in their ranging area. In non-human primates, the increase of day range, possibly with increasing group size, has been used as an index to evaluate the degree of food competition within groups. In colobines, a weak relationship between group size and day range has often been found, possibly indicating a weak food competition (Yeager and Kool 2000), though the significant relationship between the group size and ranging of the six *T. cristatus* groups likely indicates a stronger within-group feeding competition, supporting the suggestion that feeding competition in colobines may not necessarily be weak (Snaith and Chapman 2007). However, it may be difficult to conclude if the highly competitive foods in this study site are artificial or natural. We suggest that future studies on *T. cristatus* examine their food habits and nutrients as these will be key to understanding the demography in the BMKS and their socioecology in relation to intraspecific competition in folivorous primates.

Overall, we found the negative effects of provisioning on the ecology of the *T. cristatus* in the BMKS, suggesting that the study groups need effective management. Although ecotourism does not always negatively impact primate conservation (Lhota et al. 2019a; Lhota et al. 2019b), it has often failed to deliver conservation benefits for free-ranging primates, especially provisioning (Russon and Wallis 2014a). Therefore, we recommend that local authorities take appropriate actions to make visitors aware that they should stop feeding animals through educational methods explaining the negative impacts of provisioning on the ecology of *T. cristatus* in the BMKS.

## Acknowledgments

We appreciate the Department of Wildlife and National Parks Peninsular Malaysia for giving us a permit (JPHL and TN(P): 100-34/1.24 Jld 7(25) to conduct the study at BMKS. We also appreciate Kuala Selangor District Council to use their facilities at BMKS, the Metereology Department for providing our climate data, the botanist of the Department of Biological Sciences and Biotechnology, Faculty Science and Technology Universiti Kebangsaan Malaysia, Ahmad Fitri Zohari, for his help in identifying the types of trees around the study area. Finally, we are grateful to anonymous reviewers and handling editor for their fruitful comments. This study was conducted and funded under study leave scheme awarded to Norlinda Mohd-Daut by Universiti Kebangsaan Malaysia. This study was partially funded by the Society for the Promotion of Science (KAKENHI Grant Nos. #19KK0191 and # 19H03308 to IM), JSPS Core-to-Core Program, Advanced Research Networks (#JPJSCCA20170005 to S. Kohshima), and the Cooperative Research Program by KUPRI (2016-B-97).

---

**Appendix 1.**
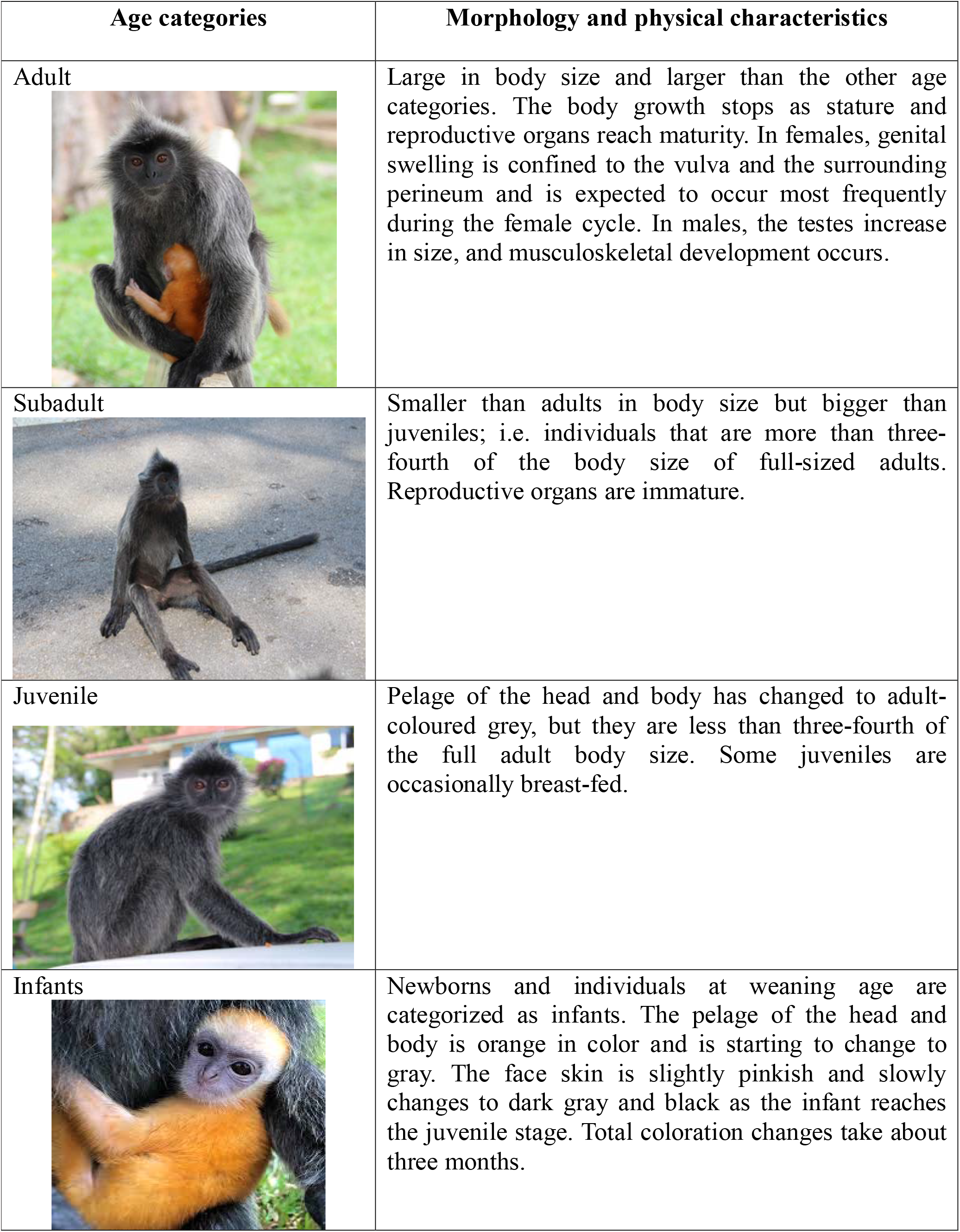
Chart of age categories with associated morphological and physical characteristics according to Aggimarangsee (2002).

